# Histologically derived fiber response functions for diffusion MRI vary across white matter fibers - an ex vivo validation study in the squirrel monkey brain

**DOI:** 10.1101/438978

**Authors:** Kurt G Schilling, Yurui Gao, Iwona Stepniewska, Vaibhav Janve, Bennett A Landman, Adam W Anderson

## Abstract

Understanding the relationship between the diffusion-weighted MRI signal and the arrangement of white matter fibers is fundamental for accurate voxel-wise reconstruction of the fiber orientation distribution (FOD) and subsequent fiber tractography. Spherical deconvolution reconstruction techniques model the diffusion signal as the convolution of the FOD with a response function which represents the signal profile of a single fiber orientation. Thus, given the signal and a fiber response function, the FOD can be estimated in every imaging voxel by deconvolution. However, the selection of the appropriate response function remains relatively un-studied, and requires further validation. In this work, using 3D histologically-defined FODs and the corresponding diffusion signal from three ex vivo squirrel monkey brains, we derive the ground truth response functions. We find that the histologically-derived response functions differ from those conventionally used. Next, we find that response functions statistically vary across brain regions, which suggests that the practice of using the same kernel throughout the brain is not optimal. Additionally, response functions vary significantly across subjects. We show that different kernels lead to different FOD reconstructions, which in turn can lead to different tractography results depending on algorithmic parameters, with large variations in the accuracy of resulting reconstructions. Together, this suggests that there is room for improvement in estimating and understanding the relationship between the diffusion signal and the underlying FOD.

## Introduction

Diffusion MRI fiber tractography has been widely adopted in a large number of clinical and basic neuroscience studies to probe the structural connectivity of the brain ^1–3^. A key requirement for anatomically correct tractography is the accurate reconstruction of the distribution of neuronal fiber orientations in each voxel, an object often referred to as the fiber orientation distribution (FOD) ^4,5^. A large number of reconstruction strategies have been proposed with the aim to estimate, model, or infer the structure of the FOD. Among them, spherical deconvolution (SD) ^6–9^ has emerged as one of the most commonly implemented in neuroscience research.

Although there are many versions and derivations of this technique ^6–20^ most spherical deconvolution methods are based on the idea that the diffusion MRI signal, S, can be modelled as the convolution of the fiber orientation distribution, with a kernel (called the fiber response function, R) that represents the signal attenuation that would be measured from a single (coherently oriented) fiber population:

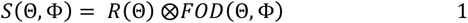

where Θ and Φ are the elevation and azimuthal angle in spherical coordinates (the response function is defined so that fibers are aligned along the vertical z-axis). The FOD can then be calculated by a deconvolution of the signal with the response function. Thus, an appropriate response function is required for accurate reconstruction of the FOD.

Several methods have been proposed to estimate the response function. Most derive the kernel from the diffusion data itself. For example, the response function can be estimated from the signal from voxels (typically 100+ voxels) with the highest fractional anisotropy values – voxels that are typically assumed to contain coherently oriented fibers ^21^. Due to relying only on FA values, this family of algorithms is susceptible to artifacts (for example, Gibbs ringing near ventricles). To overcome this limitation, several strategies have been proposed which iterate between performing SD and estimating the response function from a set of voxels most likely to contain only a single fiber population ^22,23^. Alternatively, one can assume that a single fiber bundle gives rise to a spin displacement distribution that can be described with a single diffusion tensor model, and derive the response function as the signal that would be obtained from the tensor with given shape properties ^10,14^. Despite evidence that the diffusion signal is dependent on an array of microstructural parameters (axon packing, diameter, myelination, etc.), in most cases the same response function is used in the deconvolution throughout the entire brain. There are some exemptions, where the response function varies based on tissue (i.e. white matter and gray matter) ^24,25^, however, these kernels still remain the same throughout the entire tissue type volume (for example, all white matter—an exception is the FORECAST method [8]). Despite the large number of approaches for response function estimation, little work has been done to validate the accuracy of these techniques and subsequent effects on tractography, or to question the validity of a brain-wide fiber response function.

In this study, we aim to use histology to derive the linear relationship (i.e., the response function) between the diffusion signal and the underlying FOD. Instead of the traditional approach of deconvolving the signal with an empirical response function in order to estimate the fiber distribution, we do the opposite – deconvolve the signal with the histologically-derived FOD to derive the response function. In essence, we ask, “what is the true response function, given the diffusion signal and the FOD”? To do this, we perform diffusion MRI on three fixed ex vivo squirrel monkey brains, followed by 3D confocal microscopy to derive the “ground truth” FOD. We begin by testing the validity of utilizing the same response function throughout the entire brain. Next, we assess the effects of varying response functions on both the voxel-wise reconstruction and tractography results. Finally, the anatomical accuracy of tractography using different response functions is assessed by comparing tracking results to known connections from an existing squirrel monkey atlas.

## Methods

### Overview

An overview of the study methodology is shown in Figure 1. Briefly, the ex vivo brain was first scanned using MR imaging to acquire the diffusion signal (Figure 1A; see MRI Acquisition). After sectioning, the tissue was imaged using confocal microscopy and multiple 3D confocal z-stacks were acquired (Figure 1B, see Histological Procedures). These z-stacks were processed using a pipeline previously developed to analyze 3D histology^26,27^, which resulted in voxel-wise histological ground truth 3D FODs (Figure 1C, see Histological Procedures). Because we want to estimate the response function, the diffusion signal can be deconvolved with the corresponding FODs, resulting in voxel-wise reconstructions of the response functions (Figure 1D, see Response Function Estimation). All animal procedures were approved by the Vanderbilt University Animal Care and Use Committee.

**Figure 1.**
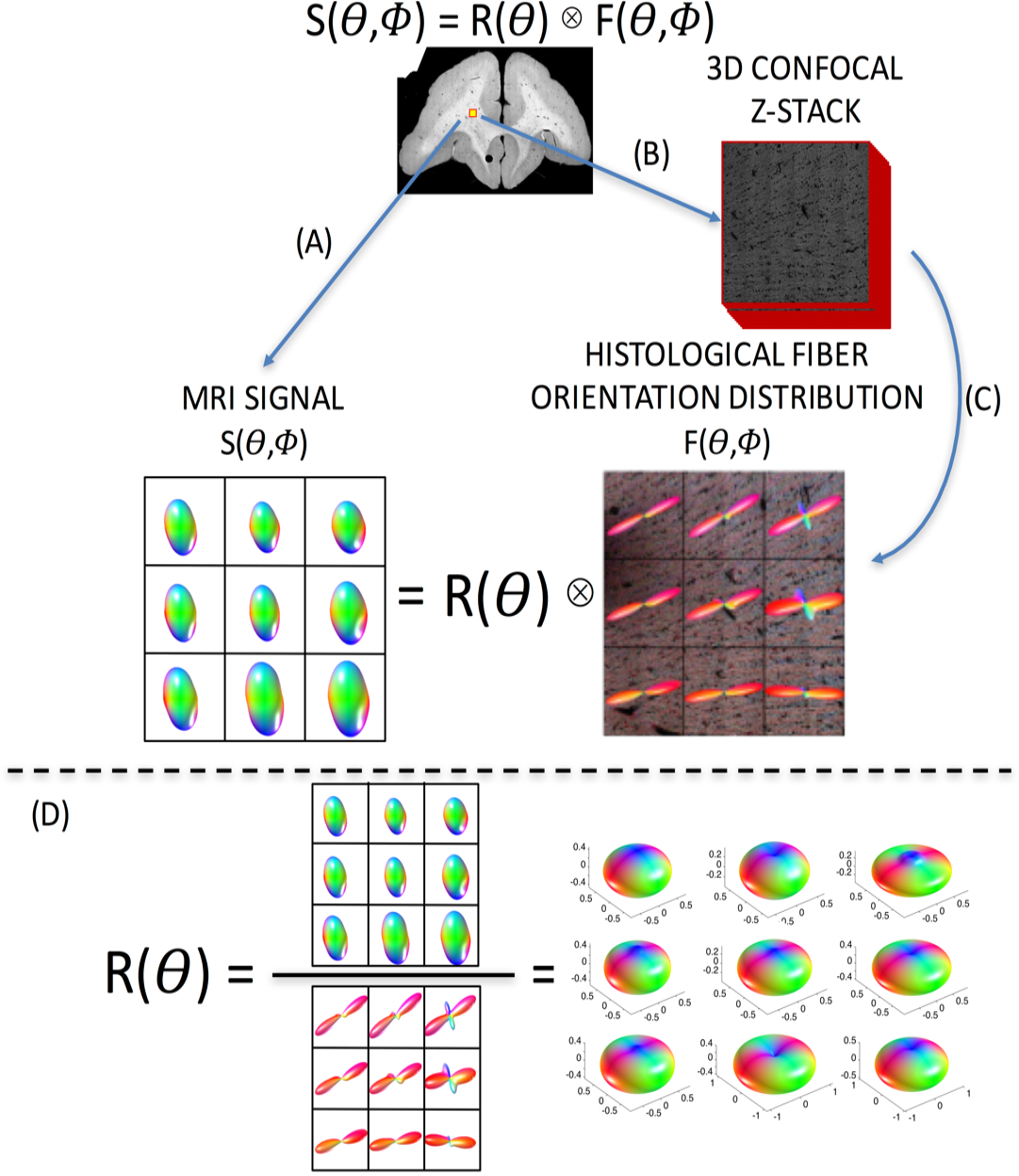
Overview of study methodology. Spherical deconvolution models the diffusion signal, S, as the convolution of the fiber response function, R, with the fiber orientation distribution, F. The brain is scanned (A) resulting in an MRI signal. Next, 3D confocal microscopy is acquired (B), followed by image processing (C), resulting in the ground truth histological FOD. Estimating the voxel-wise response function (D) is done through deconvolution of the signal with the corresponding FOD.

### MRI Acquisition

Diffusion MRI experiments were performed on three adult squirrel monkey brains that had been perfused with physiological saline followed by 4% paraformaldehyde. The brain was then immersed for 3 weeks in phosphate-buffered saline (PBS) medium with 1 mM Gd-DTPA in order to reduce longitudinal relaxation time ^28^. The brain was placed in liquid Fomblin (California Vacuum Technology, CA) and scanned on a Varian 9.4 T, 21 cm bore magnet using a quadrature birdcage volume coil (inner diameter = 63 mm).

Diffusion data were acquired with a 3D spin-echo diffusion-weighted EPI sequence (TR = 410 ms; TE = 41 ms; NSHOTS = 4; NEX = 1; Partial Fourier k-space coverage = 0.75) at 300 µm isotropic resolution. Diffusion gradient pulse duration and separation were 8 ms and 22 ms, respectively, and the b-value was set to 6000 s/mm2. This value was chosen due to the decreased diffusivity of ex vivo tissue, which is approximately a fourth of that in vivo ^29,30^, and is expected to closely replicate the signal attenuation profile for in vivo tissue with a b-value of approximately 1500 s/mm^2^. A gradient table of 107 uniformly distributed directions ^31^ was used to acquired 107 diffusion-weighted volumes with four additional image volumes collected at b = 0.

### Histological Procedures

Here, we aim to extract the histological FOD from 3D confocal z-stacks in areas equivalent to the size of an MRI voxel. To do this, we implement the methodology described in ^27^, which consisted of tissue sectioning and staining, confocal acquisition, and confocal pre-processing, followed by an image processing technique – structure tensor analysis – which resulted in an orientation estimate for every pixel in the 3D z-stack that is occupied by a fiber. The FOD is then literally the distribution of these orientation estimates. This procedure has been validated against manually traced fibers ^27^, and has been used to assess the accuracy of diffusion MRI reconstruction techniques ^26^.

Briefly, the tissue was sectioned at a thickness of 50um in the coronal plane. During sectioning, the frozen tissue block was digitally photographed prior to cutting every third section, resulting in a 3D “block-face” volume, which was used for inter-modality registration (see Image Registration).

Tissue sections were stained with the fluorescent lipophilic dye “DiI” (1, 10-dioctadecyl-3,3,30 30-tetramethylindocarbocyanine perchlorate) and imaged using an LSM 710 confocal microscope (Carl Zeiss, Inc. Thornwood, NY. USA). For all slices, acquisition began with a 2D confocal montage (0.8 × 0.8um resolution), used for image registration and for localizing the 3D high resolution region of interest. A 3D z-stack was then collected (0.2×0.2×0.42um resolution) with an in-plane field of view of 900×900um (equivalent to 9 MRI voxels). Confocal data were converted from the LSM file format to TIFF images and imported into MATLAB for image processing.

Image pre-processing consisted of attenuation correction, anisotropy correction, and interpolation to isotropic resolution. Next, pixel-wise fiber orientations in the 3D z-stack were obtained via structure tensor analysis ^32^. The histological FOD was then computed as the histogram of the extracted fiber orientations, and fit to high order (*l*_max_=20) spherical harmonic coefficients (SH). Throughout this paper, the histological-FODs are displayed as 3D glyphs in the same way that the MRI-FOD’s are typically displayed in literature.

### Image Registration

In order to derive voxel-wise response functions, it is important that the diffusion MRI signal and the histological FODs are aligned and oriented appropriately. A multi-step registration procedure 33 was used to align histology to MRI data. The first step is registration of the 2D confocal montage to the corresponding block face image using mutual information based 2D linear registration followed by 2D nonlinear registration using the adaptive bases algorithm (ABA) ^34^. Next, all block face photographs were assembled into a 3D block volume, and registered to the MRI b = 0 image using a 3D affine transformation followed by 3D nonlinear registration with ABA. Given the location of the 3D z-stack in the 2D confocal montage, we can use the combined deformation fields to determine the MRI signal from the same tissue volume. We then implemented the preservation of principle directions strategy ^35^ to the diffusion weighting vectors (b-vectors) in order to take into account rotation, scaling, and shearing effects of the spatial transformations. At this point, for each histological FOD, we have the corresponding diffusion signal – which was then fit to SH coefficients to facilitate deconvolution.

This multi-step registration procedure was validated in an earlier study ^33^, which showed that the accuracy of the overall registration was approximately one MRI voxel (~0.3 mm) by comparing position errors between manually chosen landmarks in histological and MRI space. Further, the combination of registration and histological procedures is similar to those utilized in ^27^, which showed an angular accuracy of approximately 5° in single fiber regions when compared to manually defined ground truth fiber orientations.

### Response Function Estimation

As described in Equation 1, the diffusion signal is assumed to be the convolution of the fiber distribution with the response function. Thus, to derive the response function, we *deconvolved* the signal with the histological FOD.

Following the nomenclature from ^8^, if the signal, response function, and FOD are expanded in terms of spherical harmonic coefficients s_lm_, r_lm_, and p_lm_, respectively (where l is the SH order and m is the phase), it can be shown that the deconvolution leads to a simple relationship (see Appendix A of ^8^ for complete derivation):

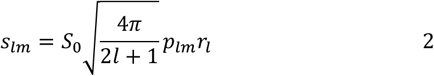

where S_0_ represents the non-diffusion weighted signal and can be assumed to be equal to 1 as long as the diffusion weighted signal is normalized to 1 before SH fitting. Inverting this gives:

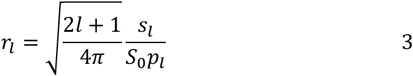

where r_l_ is only defined for m=0 (for axial symmetry of the response function around the z-axis), and p_l_ and s_l_ are vectors of coefficients (one element for each m value), for each order. Alternately, by multiplying both sides of Eq. (2) by p_lm_ and summing over m, the response function can be estimated from:

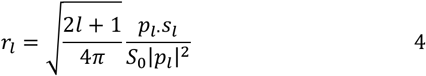

We performed this procedure for 6 z-stacks on each monkey, resulting in 18 independent confocal z-stacks, equivalent in size to 162 MRI voxels, resulting in 162 individual response function derivations. For quantitative analysis, we have chosen to focus only on voxels in the white matter which contain a single fiber population – defined to have only a single local maximum in the histological FOD. For this reason, voxels containing gray matter or crossing fibers were excluded, resulting in 153 total samples.

### Tractography and Tracking Accuracy

We next assessed the effects of various response functions on both FOD reconstruction and tractography results. As a well-studied system with known connections in the squirrel monkey brain ^36,37^, we chose to perform tractography of the motor system, with a seed region in the primary motor cortex. Both the seed region, as well as white matter and gray matter anatomical labels, are propagated from the Validate29 squirrel monkey atlas ^36^ to each individual subject’s brain.

All reconstruction and tractography was performed using the MRTrix3 software toolkit ^21^. Given a response kernel of interest, we first estimated the FOD from diffusion data using spherical deconvolution ^7^ (command dwi2fod). From a field of FOD estimates, we performed tractography (iFOD2 algorithm) using the primary motor cortex seed as a region of interest.

The anatomical accuracy of tractography connections to labelled regions of interest was assessed by comparing results to known connections derived from histology and defined in the atlas ^36,37^. We measured the sensitivity (the ability to detect true connections) and specificity (the ability to avoid false connections) of the various tractographic implementations derived using different response kernels.

## Results

### Histologically Derived Average Response Functions

Response functions derived using standard practice methods for estimation are visualized in Figure 2 for all three subjects (A-C). Estimation using the signal from the voxels with the highest FA ^21^ (“FA Response”), using an iterative estimation method ^23^ (“Iterative Response”), and assuming a cylindrically symmetric diffusion tensor model ^10^ (“Tensor Response”) show the standard, expected, oblate shape, where the signal is lowest in the direction of fibers (along the z-axis) and highest perpendicular to them. Qualitatively, there are differences in the “flatness” of these functions within the same brain, suggesting fairly large variation in current implementations of these techniques.

**Figure 2.**
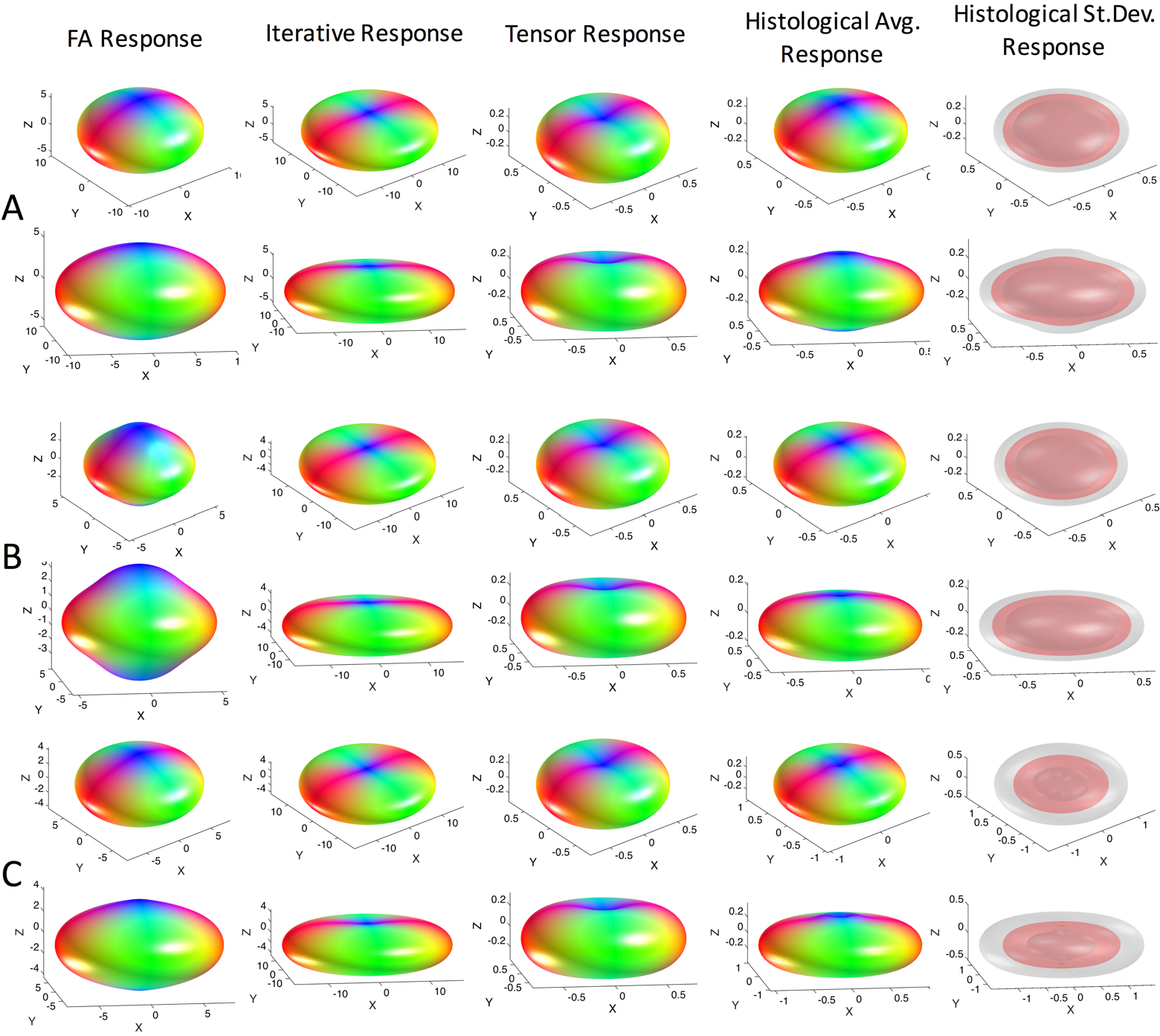
Response functions estimated using conventional methods differ from the histologically-derived response functions. Kernels estimated using the highest FA voxels, the iterative calibration method, and modelled from the diffusion tensor are shown in two views. The average histologically-derived response function for each monkey is shown (monkeys #1-3 correspond to A-C) along with the mean (red) and mean +/− one standard deviation (gray) surfaces to highlight variability. Note differences in magnitude are due to differing normalization in our work and in MRTrix3. Shape, and not magnitude, determines the final FOD shape.

The average response function across all histology is shown in Figure 2 for each subject. Qualitatively, in all cases, the histologically-derived average response functions share similarities to standard-practice methods, and are somewhat intermediate in shape between the iterative estimation methods and the tensor and FA-derived shapes. Finally, visualizing the mean and standard deviation across the brain suggests that there is some deviation in response size within white matter.

### Variation across the brain

The variation in histological response leads us to ask whether the use of a single response function across the entire brain is appropriate. For each monkey, we visualize the results of three (of the six) z-stacks (Figures 3, 4, and 5, for monkeys 1, 2, and 3, respectively), showing the z-stack location, the voxel-wise histological FOD, diffusion signal, and derived response functions, along with two views of the average response function at each location. Differences in shape, flatness, and radial and axial attenuations are apparent both within and across monkeys.

**Figure 3.**
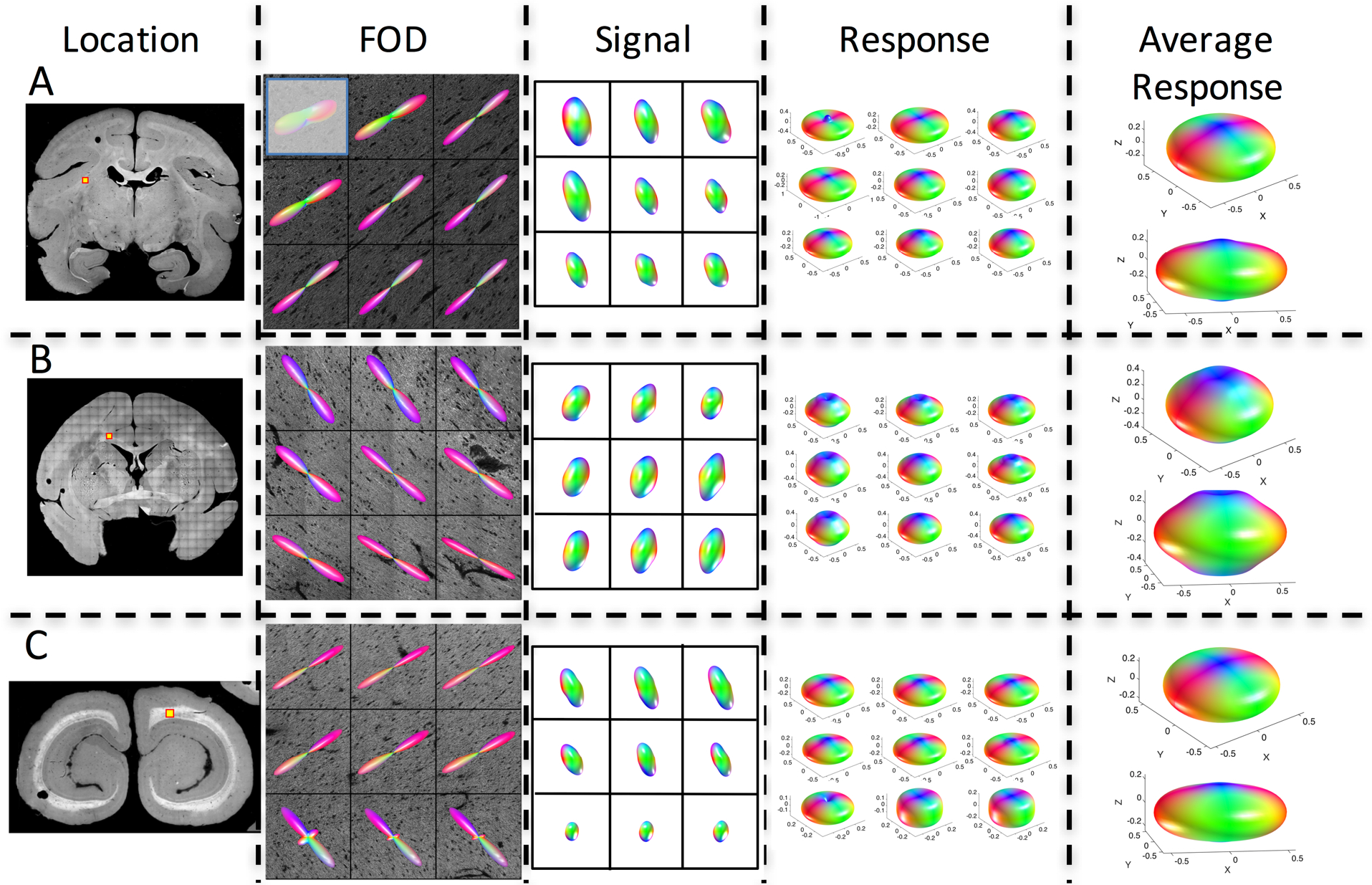
The response function varies across brain regions. Three locations in the same monkey brain (monkey #1) are shown, along with the histological fiber orientation distribution, signal, and derived response function for all 9 voxels. In addition, the average response function for each z-stack is shown in two views. Grayed-out voxel contained some gray matter and hence was not included in the study.

**Figure 4.**
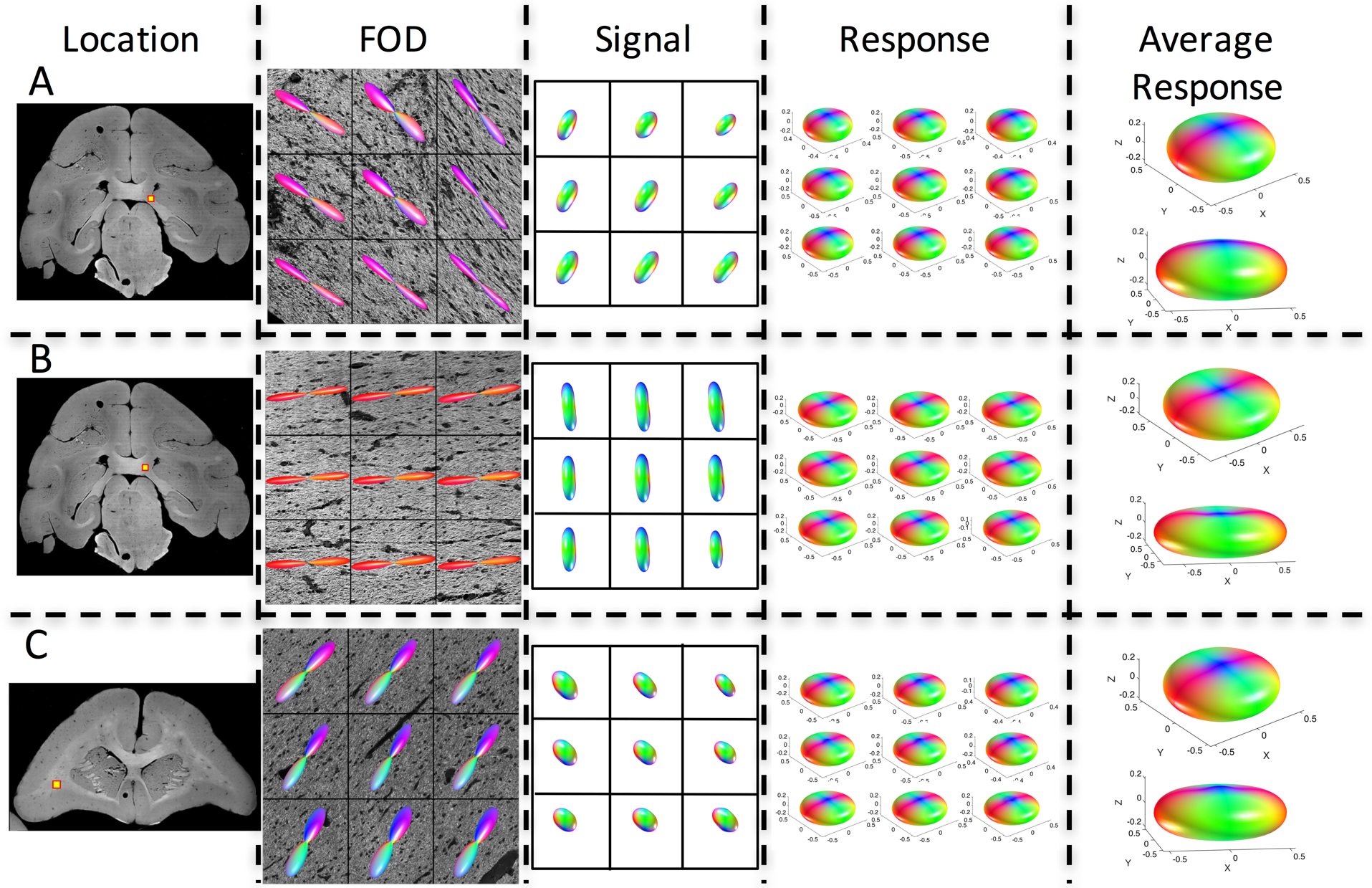
The response function varies across brain regions. Three locations in the same monkey brain (monkey #2) are shown, along with the histological fiber orientation distribution, signal, and derived response function for all 9 voxels. In addition, the average response function for each z-stack is shown in two views.

**Figure 5.**
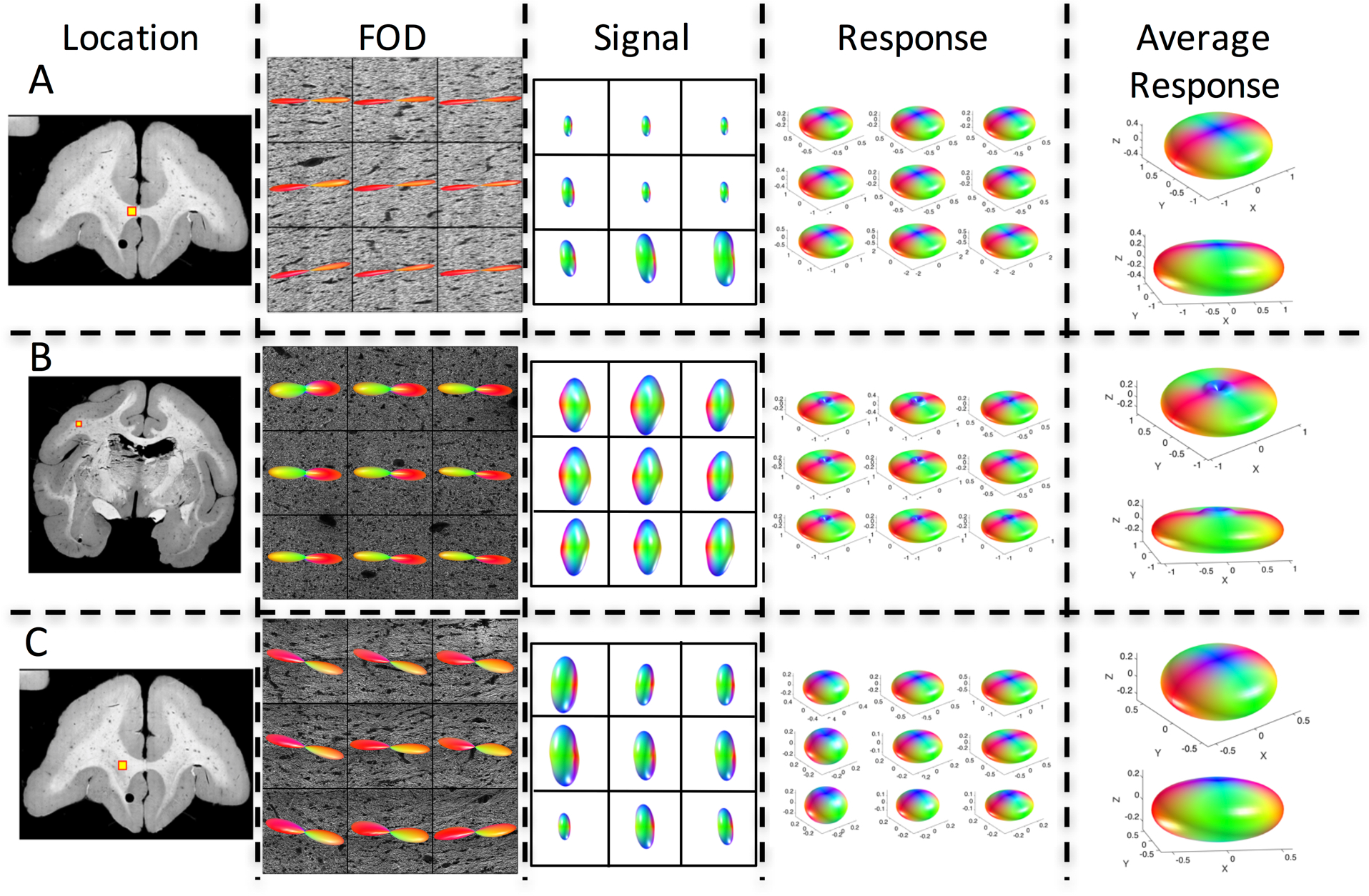
The response function varies across brain regions. Three locations in the same monkey brain (monkey #3) are shown, along with the histological fiber orientation distribution, signal, and derived response function for all 9 voxels. In addition, the average response function for each z-stack is shown in two views.

A multivariate analysis of variance (MANOVA) of the (n=4) spherical harmonic coefficients representing the response functions indicates that the response functions vary significantly across white matter locations within a brain (monkey #1: dimensionality = 3, p<.001; monkey #2: dimensionality = 3, p<.001; monkey #4: dimensionality = 4, p<.001 – here dimensionality is an estimate of the dimension of the space spanned by the group means, if the response functions were the same, the dimension would be 0). Additionally, a MANOVA also indicates that the response functions show significant differences across brains (dimensionality = 2; p<.001, indicating that the mean response function is not the same).

### Fiber orientation distribution estimation

The standard-practice kernels, as well as three histologically defined kernels from the same monkey (Monkey #1 from Figure 3) are used to perform spherical deconvolution, and the resulting FODs are shown in Figure 6 for two white matter locations. It is clear that different response functions lead to different FODs. The most obvious differences are the amplification of smaller peaks in some of the methods utilizing the “flatter” kernels (for example, the Iterative Response). Our results are in agreement with literature ^7,8^, where different response functions result in FODs with peaks in the same orientations, and only differences in relative volume fractions of different fiber populations. Given that the peak orientations are the same, the question then becomes whether this affects subsequent tractography.

**Figure 6.**
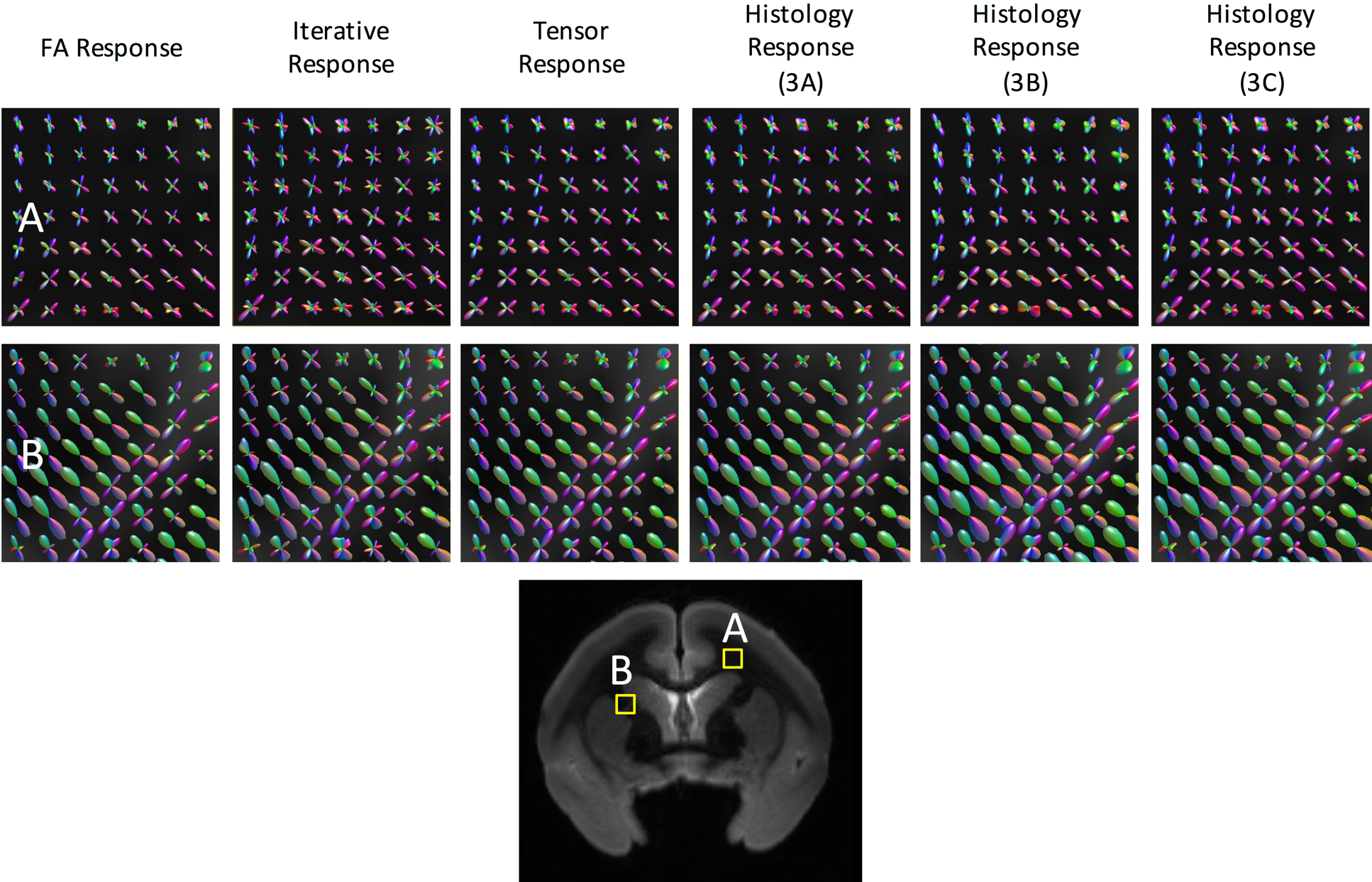
Different response functions lead to different FOD reconstructions. Fiber reconstruction in two regions of interest (A and B) using three commonly implemented response functions (FA Response, Iterative Response, Tensor Response), and three histologically derived response functions (those from Figure 3A, 3B, and 3C) from the same monkey. Reconstructed fiber orientation distributions retain similar shape and orientations, but contain different volume fractions of fiber populations.

### Tractography

Tractography from a region of the primary motor cortex (Figure 7A) was performed using the FOD reconstructions from the standard kernels, and the individual kernels from three z-stacks (the same from Figure 6). Differences in spatial extent and path representation are apparent across different response functions (Figure 7B).

**Figure 7.**
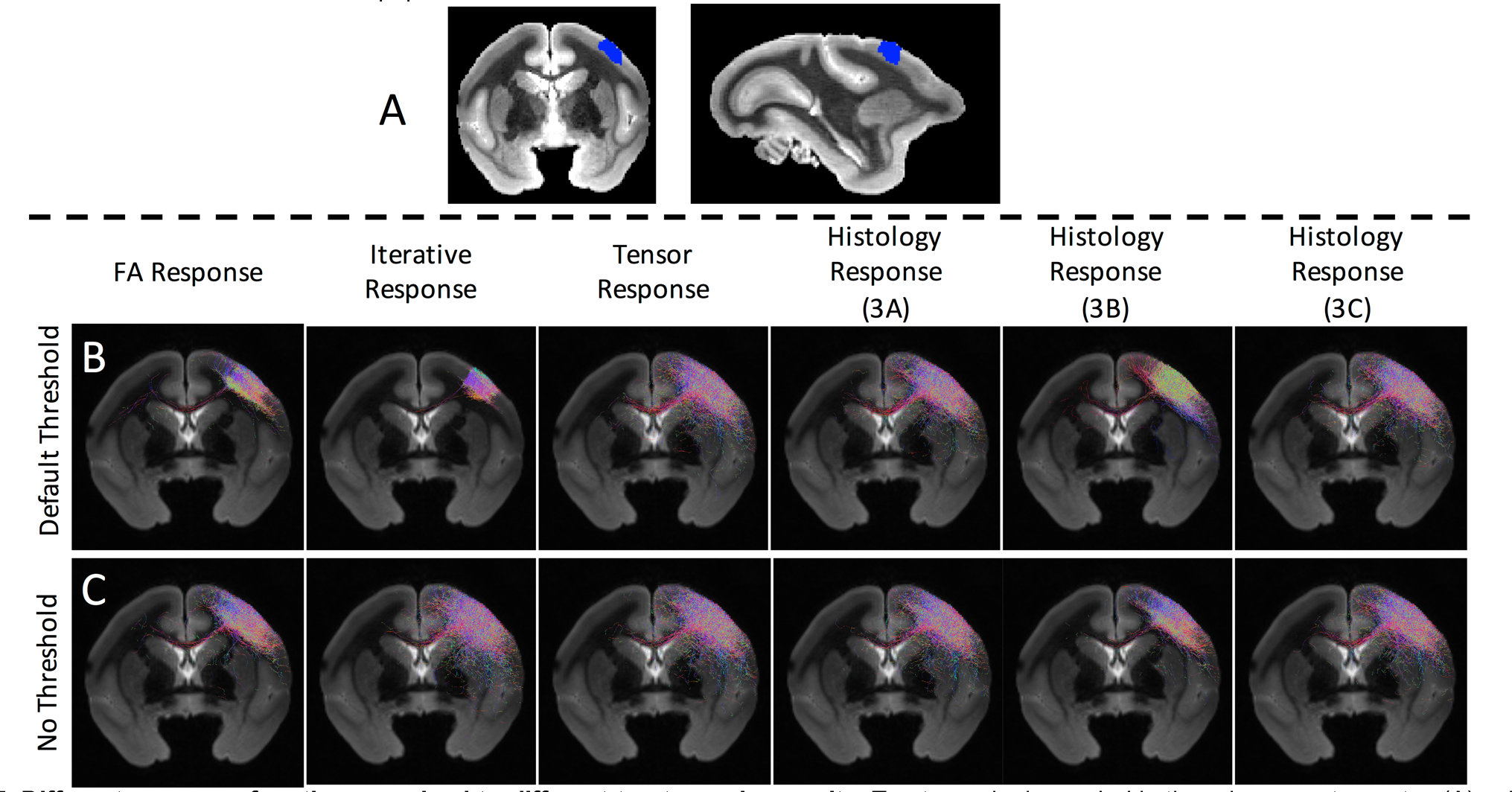
Different response functions *can* lead to different tractography results. Tractography is seeded in the primary motor cortex (A) using the FODs described in Figure 6, using standard tractography parameters and settings (i.e., the FOD amplitude threshold) (B). Tractography is repeated using no threshold on FOD amplitude, resulting in qualitatively similar streamlines (C).

Because the only differences in reconstructed FODs are the amplitude of the peaks (and not orientation), we hypothesized that differences in tractographic reconstructions are due to the amplitude threshold on the FOD used in the fiber tracking algorithms. Given a number of “peaks” or local maxima of the FOD, the “FOD threshold” determines an amplitude below which the peak is considered negligible, and is not used to propagate streamlines. For Figure 7B, this threshold was set at a typical value of 0.05 of the maximum peak amplitude. To investigate whether the threshold affected tractography, we set the threshold to zero (no threshold on peak amplitude) and re-ran tractography (Figure 7C). Qualitatively, reconstructions appear much more similar despite differences in response functions.

Connectivity to the motor cortex is compared to known connections from tracer studies. Of 71 white and gray matter ROIs (Figure 8A) propagated from the atlas, 44 are known to be connected to the left seed region, and 27 are not connected. Sensitivity and specificity of all tracking strategies from Figure 7 are computed and shown in Figure 8B for standard tracking parameters, and Figure 8C for the non-thresholded tracking. With default tracking parameters, not only do different response functions lead to qualitatively different tractography results, they can lead to a fairly wide variation in the true positive rate and true negative rate of tractography connections (Figure 8B). However, this variation is largely caused by the amplitude threshold, and when eliminated, results in very similar sensitivity/specificity results (Figure 8C).

**Figure 8.**
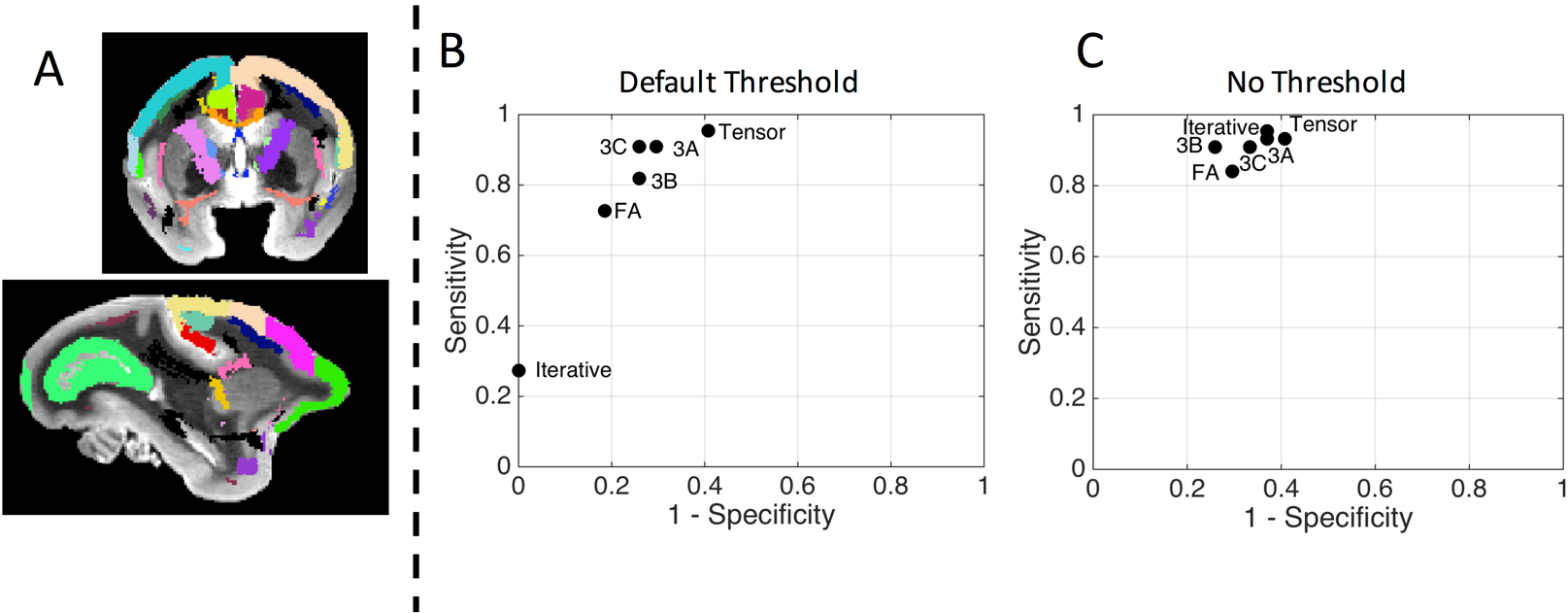
Different response functions *can* lead to different tractography results. Tractography is compared to an atlas of known connections, by calculating the true positive and true negative rate of connections to various regions of interest (A). The sensitivity and specificity of the tractography results from Figure 7 are plotted in an ROC curve using the default FOD threshold (B) and no FOD threshold (C). Method abbreviations: FA: average FA response; ITER: iterative response; Tensor: tensor-derived response; 3A-C: histological-derived response from z-stacks A-C from Figure 3.

## Discussion

Understanding the relationship between the diffusion MRI signal and tissue microstructure is a fundamental goal in the field of diffusion MR imaging. Here, we study one aspect of the microstructure, the geometric distribution of fibers in each imaging voxel (i.e., the FOD), a function that is fundamental in creating maps of the human brain using tractography. In this work, we use histology to derive the linear relationship between the diffusion signal and the underlying FOD – the response function (Figure 1).

We first observe that the histologically-derived response functions can differ in shape from conventional methods of estimating this kernel from the diffusion data itself or by assuming signal profiles (Figure 2). Differences are critical, because tractography results not only depend on tracking parameters, algorithms, and constraints (i.e. step size, curvature threshold, FA threshold) ^38–40^, but spherical deconvolution results depend on the derivation and shape of the response function. Next, we show that the histologically derived response function varies within brains and between brains (Figure 3-5). Variation across white matter is expected, as the diffusion process is known to be dependent on local anatomy and microstructural properties. Despite this knowledge, a brain-wide response function is still the most prevalent method of performing SD today. Our results demonstrate that this practice is likely not adequate for accurate reconstruction of FODs across the brain, especially with varying tissue properties. It would be useful to extract other microstructural measures, for example axon diameters, volume fraction, or fiber dispersion, possibly from the confocal data itself, to study the response functions dependencies on the microstructure.

Unsurprisingly, different response functions lead to different FOD reconstruction estimates (Figure 6). Despite differences in the apparent volume fractions of fiber bundles, the peak orientation remains the same (given that the response functions used are axially symmetric). Because of this, it was surprising that different response functions, can lead to different tractography results (Figures 7 and 8). This variation in tractography results is almost certainly dependent on the algorithmic parameters as well; for example, the threshold below which peaks are deemed “spurious” as well as how the propagation steps are determined.

To confirm this, tractography with no threshold resulted in very similar qualitative and quantitative results for all response functions, and subsequent FOD estimations. While different choices of tracking parameters can produce large variations, it is important to point out that, with typical usage (i.e., typical parameters), different response functions also lead to different tractography results.

It is also important to point out that there are a large number of spherical deconvolution methods in literature ^6–9,11–20^, each with slightly different derivations, modifications, or assumptions on the response kernels and models ^41,42^. Here, we have chosen to assess what we believe to be the most commonly utilized in the diffusion literature, implemented using common or open-sourced platforms, and do not intend to compare or rank the different methodologies, instead highlighting a difference between these and histologically derived kernels. More importantly, with histological analysis alone, we find that the response functions vary across different white matter regions, and can potentially (depending upon tracking parameters) result in different tractography reconstructions, as well as reconstructions different from existing reconstruction methods. This does not mean that the histologically informed tractography is actually “better”. In fact, the histological response may only be appropriate for FOD reconstruction in certain anatomical locations and not others, and no method was perfect (Figure 8), all exhibiting a sensitivity/specificity tradeoff previously described in the validation literature ^38–40,43,44^. The tractographic analysis serves to emphasize that different response functions can lead to different connectivity results.

### Limitations

This study has several potential limitations. First, because of the ex vivo nature of the tissue, and reduced ex vivo diffusivity ^30^, the derived response function is not the response function that should be used for in vivo data (or human data). However, studying ex vivo tissue is the only way in which the true fiber orientation distribution can be obtained, and our goal is simply to determine the linear relationship between this and the signal, and compare these data-driven methods of estimating the response function. Although these subjects did not have an in vivo acquisition, future studies should include both in vivo and ex vivo imaging, as well as multiple diffusion shells or diffusion times, followed by histological analysis, in order to better understand the response function, the signal, and the fiber distribution. Because of the ex vivo nature of the study, our results may not hold for in vivo human studies, although we expect the response function variation across brain regions, and across brains, to be largely generalizable to humans.

The second limitation is that only one connection was tested to assess differences in tractography. The primary motor system in the non-human primate has been studied extensively ^45,46^ with well characterized connections. Our approach highlights the effects of the response function on tracking accuracy, and the resulting tradeoff in tractography sensitivity and specificity is likely to exist in all pathways of interest. However, the focus of the study is on variation in the response function themselves, rather than a thorough validation of tractography ^43^.

Most importantly, for these results to be valid, it is important that the FODs derived from histology can actually be considered ground truth. The pipeline to extract 3D FODs from 3D confocal z-stacks is described thoroughly in ^27^, where the technique was validated to ensure directional accuracy on the order of 5°. Importantly, this pipeline both thresholds the image to ensure that orientation estimates are extracted only from voxels occupied by fibers, excluding directions estimated from non-tissue pixels, and also excludes voxels with poor, or uncertain, directional information. In this way, we excluded extra-axonal compartments in the FOD estimate, and FOD estimates are weighted by the volume fraction of the axons (i.e. big axons have more fiber-occupying voxels than small axons), in the same way that dMRI is sensitive to the volumes of various compartments.

Estimates of the response function can be corrupted by two sources, noise and misalignment. We do expect some misalignment of the FOD and signal due to tissue distortions and registration errors, as well as noise in the MRI signal – our ex vivo acquisitions resulted in an SNR of the b0 images in the WM of approximately 25, 29, and 22 for monkeys #1-3, typical of that in research studies. Figure 9 shows the results of simulations investigating both misalignment and noise. In this case, with uncorrupted data (SNR=∞) misalignment between FOD and signal results in very little change to the derived response function (decreased radial and increased axial signal) with as much as 20° of misalignment. In addition, with SNR as low as 10, the overall shape of the response function is qualitatively in agreement with the true function – noise causes a “blurring” of the true FOD, again decreasing radial signal and increasing axial signal. Additionally, spurious peaks frequently occurred axially in the direction where lowest signal is expected. Further analysis indicated that this peak occurs when noise is specifically along, or near, the peak direction of the FOD, and becomes more prominent as the noise level in that direction increases (Supplementary Figure 1). Noise in other directions is more regularized due to the increased density of sampling at varying azimuthal angles (i.e., longitudinal planes). In our data, these peaks were infrequent (for example, this could be observed in Figure 3A), and if they occurred, were not as prominent as in simulations.

**Figure 9.**
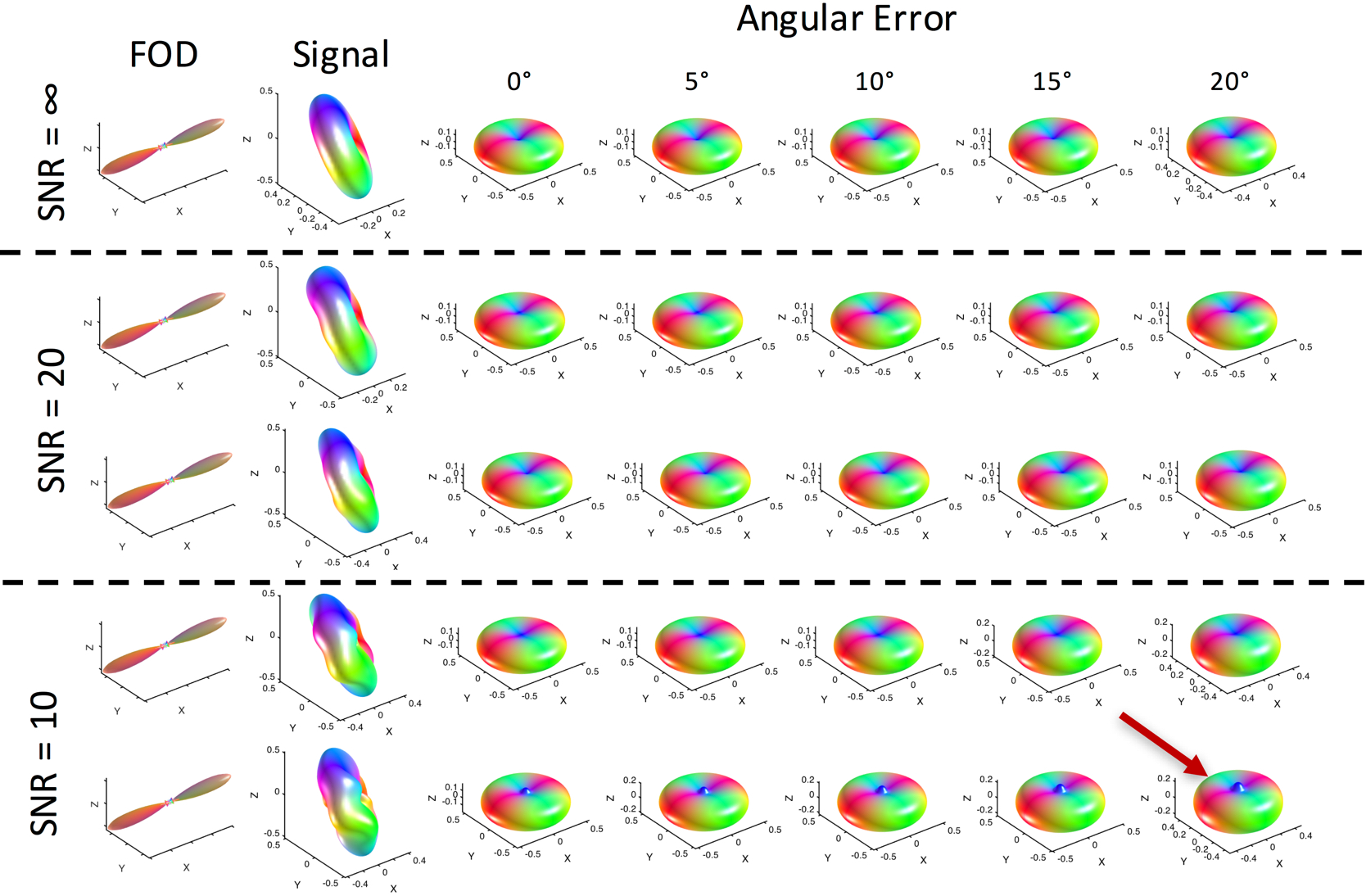
Noise and misalignment can cause errors in estimating the response functions in single fiber regions. FODs and signal at varying noise levels (SNR = ∞, 10, and 20) and misalignments (angular errors of 0-20 degrees) were used to derive response functions. Response functions are relatively robust to angular error, but noise can cause spurious peaks axially (red arrow). Here, signal is generated from an axially symmetric diffusion tensor model with λ_1_ = 1.6E-3 mm^2^/s, λ_2_ = λ_3_=1.6E-3 mm^2^/s, FA=0.7. Two simulations are shown for each noise level. Noise is added in quadrature.

### Crossing Fibers

This study focused on single fiber voxels in the white matter, in order to avoid contamination or partial volume effects from other compartments – including gray matter or multiple intra-voxel fiber populations. However, the dataset collected in ^26^ additionally contains a number of z-stacks that contain crossing fibers (defined as voxels having multiple local maxima in the FOD). Figure 10 shows example z-stacks with crossing fibers (which were not used in the current study), highlighting location, shape of the FOD shown in two views (to highlight crossing and show both in-plane and through-plane fiber orientations), and the derived response function. We also show RGB colormaps (Figure 10D) for visualizing crossing fiber patterns.

**Figure 10.**
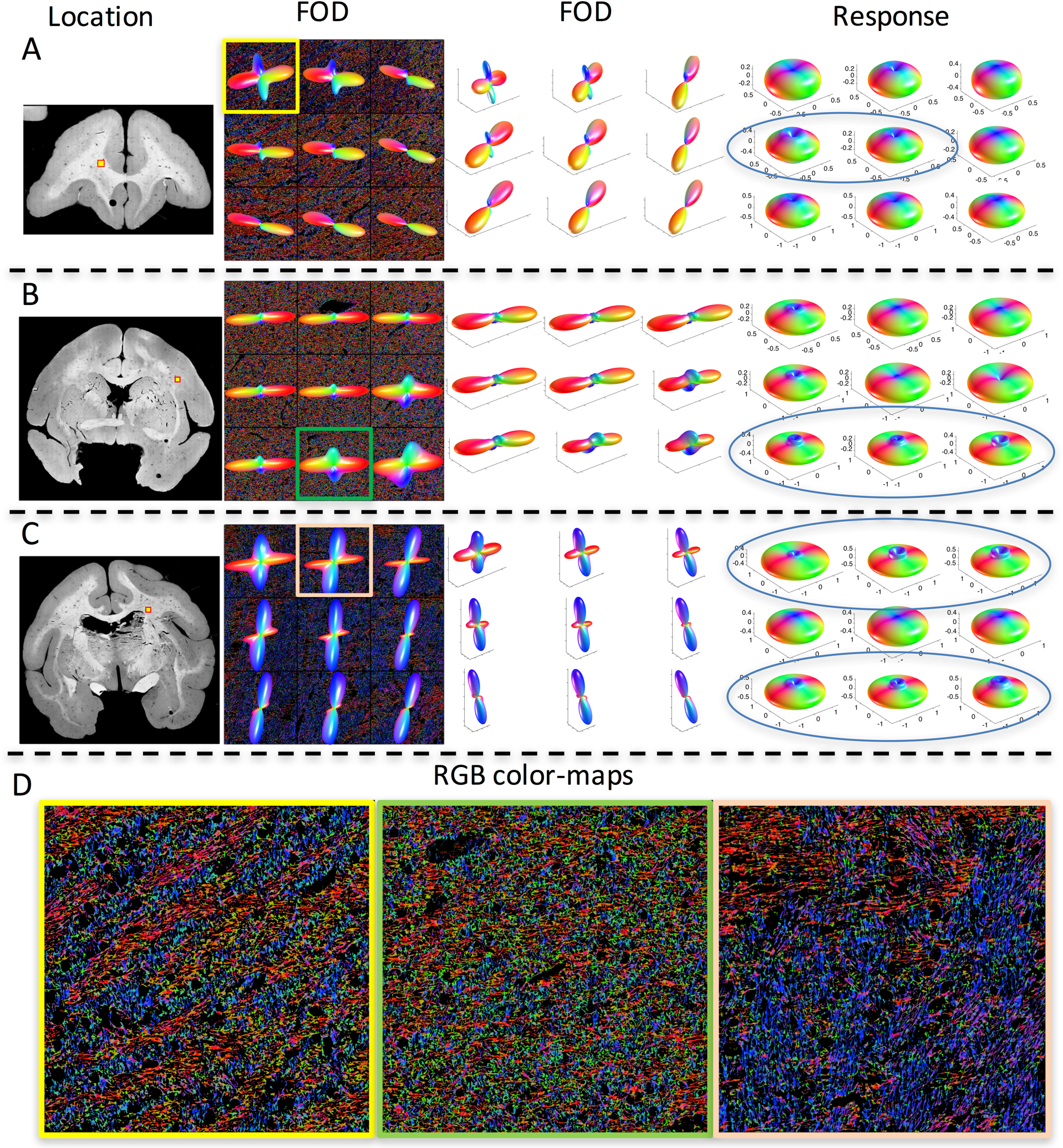
Histologically-derived response functions in crossing fiber regions display unconventional shapes. Three locations determined to contain crossing fiber populations (A-C) are shown, along with the FOD (viewed along the normal to the tissue section) overlaid on RGB color-maps, the FOD shown at an alternate angle (to highlight both in-plane and through plane fibers), and the derived response functions. Many response functions show a “cusp-like” shape axially (indicated by blue ellipses). RGB color-maps of individual voxels (D) highlight orientation estimates (Red: right/left, Green: anterior/posterior, Blue: superior/inferior) of every pixel in the 3d z-stack volume occupied by a fiber and show complex, interwoven, crossing fiber patterns.

An interesting observation across many of the crossing fiber regions is the presence of a “cusp-like” shape axially, instead of a simple monotonically decreasing function at decreasing polar angles (towards z-axis). There are several potential explanations. First, this could be a genuine effect of the diffusion process due to boundaries and restrictions, especially at angles close to (within 5-20 degrees) of the main fiber orientation. This could be caused by a combination of multiple diffusion pools (for example, one or more for each fiber population), where the process cannot be adequately explained by a single axial and radial diffusivity. Alternatively, this could be an artifact. Investigating the effects of noise and misalignment through simulations on crossing fibers (Figure 11) shows that noise can, indeed, cause a cusp-like shape (red arrow), in addition to the spurious peaks directly along the azimuth (blue arrows). However, this generally happened at an SNR lower than in our acquired data, and was relatively infrequently observed in simulations. While we found that this “cusp” did in fact appear in simulations when two *different* response functions exist within an imaging voxel (Figure 11; also see Supplementary Figure 2 for 100 response function estimations at an SNR=10 when two different response functions exist), the cusp did not appear in simulations when the *same* response function describes both fiber populations (see Supplementary Figure 2 for 100 estimations at an SNR=10 when one response function describes both fiber populations). When the signal is generated with two fibers with inherently different response kernels, we believe that the deconvolution procedure is less stable, and can result in erroneous high order coefficient estimates (giving rise to the sharp cusp), and an overall estimate intermediate between the two true kernels. Whereas with the same response function, the estimate is stable, and noise produces a uniform blurring of the estimate (as in single fiber populations). Although the focus of this study is not on crossing fibers, we believe that these simulations suggest that in addition to variation within and across brains, response functions can vary within voxels. Additionally, the cusp shape was seen (although only in one z-stack, Figure 5B) in single fiber populations, which may suggest that there are indeed two fiber populations with two different response functions, which share the same or similar orientation (a phenomenon expected in the brain ^44^), although only one local maximum is identified in the FOD. Again, the possibly supports the argument that response functions can vary within a voxel.

**Figure 11.**
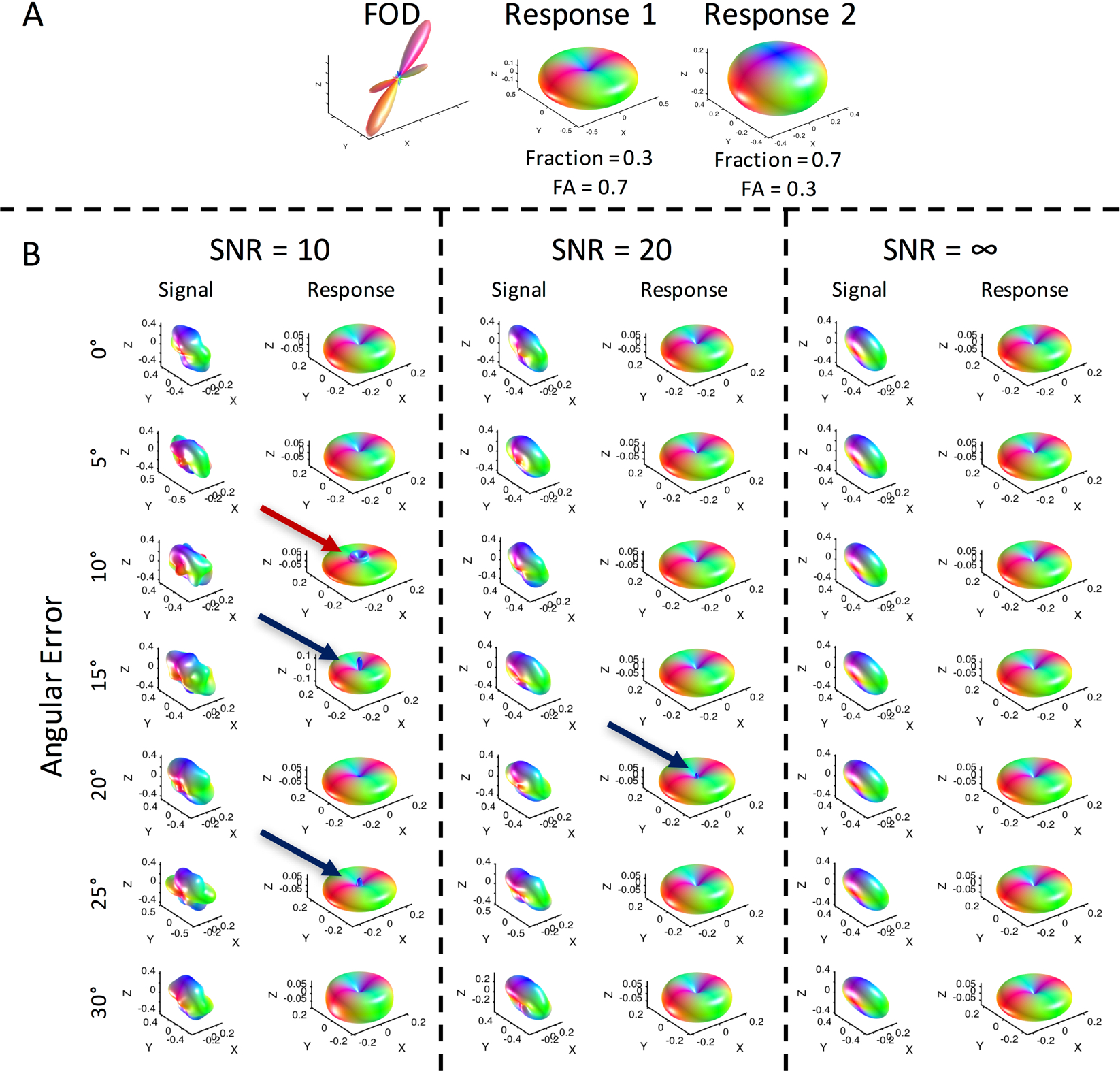
Noise and misalignment can cause errors in estimating the response functions in crossing fiber regions. FODs and signal at varying noise levels (SNR = ∞, 10, and 20) and misalignments (angular errors of 0-30 degrees) were used to derive response functions. Noise can cause spurious peaks axially (blue arrows), and can result in “cusp-like” shapes near the poles.

## Conclusion

Here, for the first time, we have empirically estimated the linear relationship (i.e., the fiber response function) between the diffusion signal and the underlying FOD using 3D histological analysis. We find statistically significant differences in response functions between brain regions, which suggests that a single response function may not be appropriate for use throughout the entire brain, and instead a region-wise, or fiber-wise, response may be more appropriate. Additionally, we find different response functions across different brains, as well as evidence that response functions may vary within voxels due to multiple fiber populations. Differences in response functions lead to different FOD reconstructions, which in turn can lead to different tractography results, even though peaks in the FOD remain the same orientation.

## Acknowledgments

This work was supported by the National Institute of Neurological Disorders and Stroke of the National Institutes of Health under award numbers RO1 NS058639 and S10 RR17799. Whole slide imaging was performed in the Digital Histology Shared Resource at Vanderbilt University Medical Center (www.mc.vanderbilt.edu/dhsr). We would also like to thank reviewers for recommendations on analysis and on deriving the response function (Eq. 4).

---

**Supplementary Figure 1.**
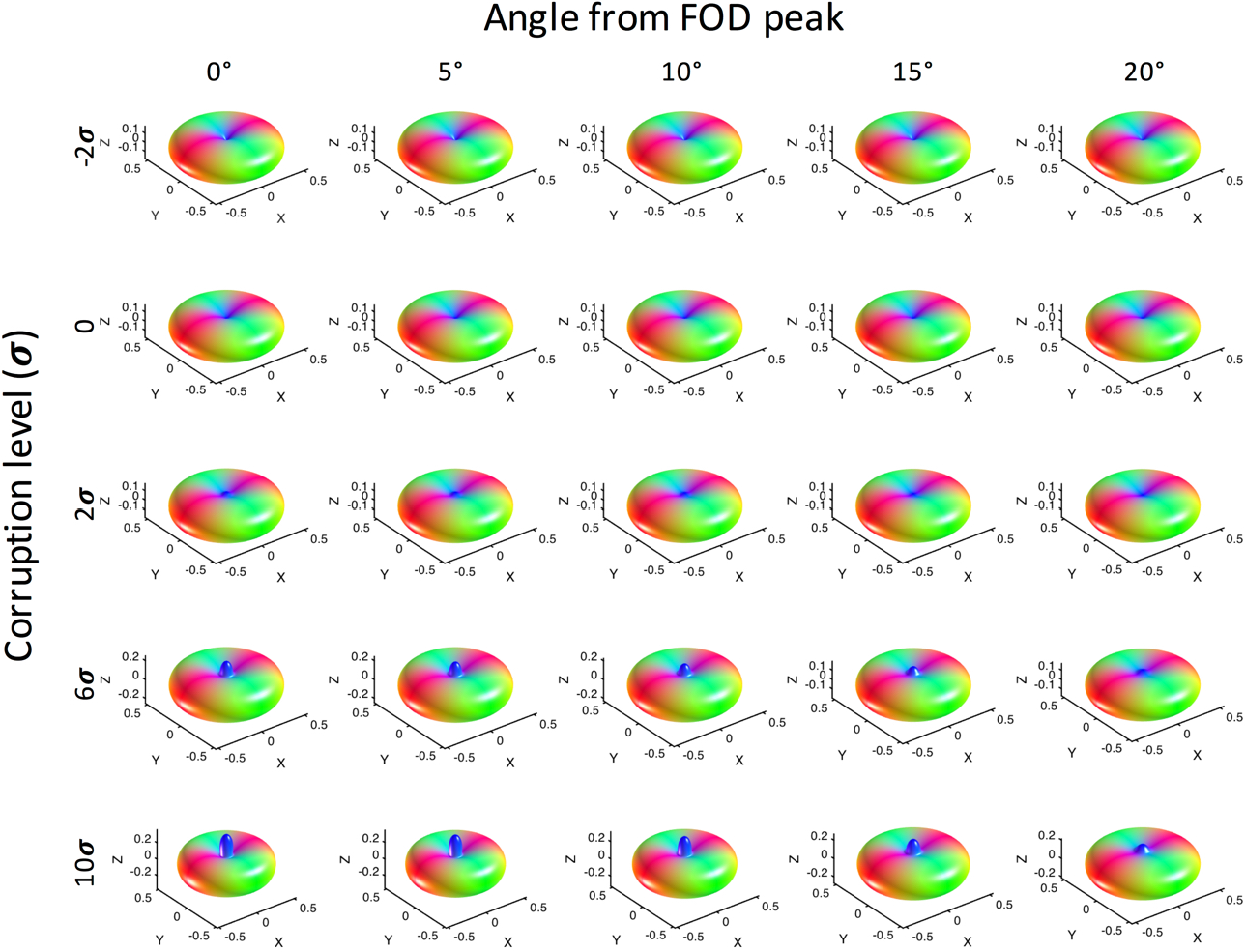
Noise in the direction of the FOD peak causes axial spikes in the estimated response function. Signal was corrupted with varying levels of noise (−2 to 10 standard deviations) along the direction of the FOD peak (minimum signal) and at varying angles from the peak (0 to 20 degrees off axis).

**Supplementary Figure 2.**
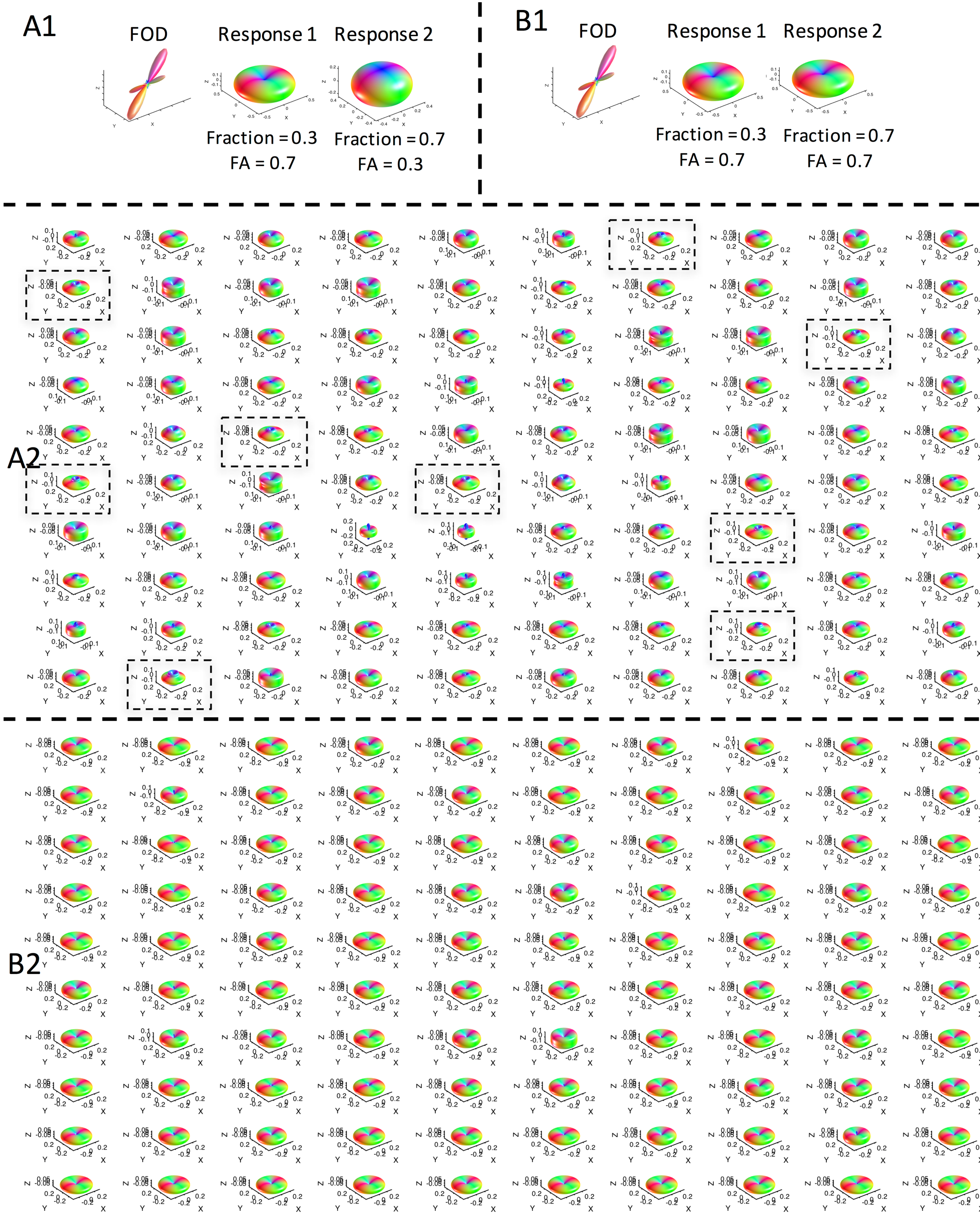
Cusps in the estimated response function only appear if the true signal is composed of 2 different true response functions. With two different response functions (A1), and an SNR of 10, cusps in the estimated response function appear frequently (A2, black boxes). Each glyph represents a separate simulation in the presence of noise. However, with the same response function (B1), and an SNR of 10, no cusps are apparent in the estimated response function.

## References

1. Johansen-Berg H, Behrens TE. Just pretty pictures? What diffusion tractography can add in clinical neuroscience. Curr Opin Neurol. 2006;19(4):379–385.

2. Catani M, Thiebaut de Schotten M. Atlas of human brain connections. 2015.

3. Jones DK. Diffusion MRI : theory, methods, and application. Oxford; New York: Oxford University Press; 2010.

4. Alexander DC, Seunarine KK. Mathematics of crossing fibers. In: Jones DK, ed. Diffusion MRI: theory, methods, and application. Oxford; New York: Oxford University Press; 2010:451–464.

5. Frank LR. Characterization of anisotropy in high angular resolution diffusion-weighted MRI. Magnetic resonance in medicine : official journal of the Society of Magnetic Resonance in Medicine / Society of Magnetic Resonance in Medicine. 2002;47(6):1083–1099.

6. Tournier JD, Calamante F, Connelly A. Robust determination of the fibre orientation distribution in diffusion MRI: non-negativity constrained super-resolved spherical deconvolution. NeuroImage. 2007;35(4):1459–1472.

7. Tournier JD, Calamante F, Gadian DG, Connelly A. Direct estimation of the fiber orientation density function from diffusion-weighted MRI data using spherical deconvolution. NeuroImage. 2004;23(3):1176–1185.

8. Anderson AW. Measurement of fiber orientation distributions using high angular resolution diffusion imaging. Magnetic resonance in medicine : official journal of the Society of Magnetic Resonance in Medicine / Society of Magnetic Resonance in Medicine. 2005;54(5):1194–1206.

9. Anderson AW, Ding Z. Sub-voxel measurement of fiber orientation using high angular resolution diffusion tensor imaging. Paper presented at: Tenth Scientific Meeting of the International Society for Magnetic Resonance in Medicine 2002.

10. Canales-Rodriguez EJ, Legarreta JH, Pizzolato M, et al. Sparse wars: A survey and comparative study of spherical deconvolution algorithms for diffusion MRI. NeuroImage. 2018;184:140–160.

11. Canales-Rodriguez EJ, Daducci A, Sotiropoulos SN, et al. Spherical Deconvolution of Multichannel Diffusion MRI Data with Non-Gaussian Noise Models and Spatial Regularization. PloS one. 2015;10(10):e0138910.

12. Cheng J, Deriche R, Jiang T, Shen D, Yap PT. Non-Negative Spherical Deconvolution (NNSD) for estimation of fiber Orientation Distribution Function in single-/multi-shell diffusion MRI. NeuroImage. 2014;101:750–764.

13. Daducci A, Van De Ville D, Thiran JP, Wiaux Y. Sparse regularization for fiber ODF reconstruction: from the suboptimality of l2 and l1 priors to l0. Med Image Anal. 2014;18(6):820–833.

14. Dell’acqua F, Scifo P, Rizzo G, et al. A modified damped Richardson-Lucy algorithm to reduce isotropic background effects in spherical deconvolution. NeuroImage. 2010;49(2):1446–1458.

15. Kaden E, Kruggel F. Nonparametric Bayesian inference of the fiber orientation distribution from diffusion-weighted MR images. Med Image Anal. 2012;16(4):876–888.

16. Kaden E, Knosche TR, Anwander A. Parametric spherical deconvolution: inferring anatomical connectivity using diffusion MR imaging. NeuroImage. 2007;37(2):474–488.

17. Sakaie KE, Lowe MJ. An objective method for regularization of fiber orientation distributions derived from diffusion-weighted MRI. NeuroImage. 2007;34(1):169–176.

18. Scherrer B, Schwartzman A, Taquet M, Sahin M, Prabhu SP, Warfield SK. Characterizing brain tissue by assessment of the distribution of anisotropic microstructural environments in diffusion-compartment imaging (DIAMOND). Magnetic resonance in medicine : official journal of the Society of Magnetic Resonance in Medicine / Society of Magnetic Resonance in Medicine. 2016;76(3):963–977.

19. Sotiropoulos SN, Behrens TE, Jbabdi S. Ball and rackets: Inferring fiber fanning from diffusion-weighted MRI. NeuroImage. 2012;60(2):1412–1425.

20. Tristan-Vega A, Westin CF. Probabilistic ODF estimation from reduced HARDI data with sparse regularization. Med Image Comput Comput Assist Interv. 2011;14(Pt 2):182–190.

21. Tournier JD, Calamante F, Connelly A. MRtrix: Diffusion tractography in crossing fiber regions. International Journal of Imaging Systems and Technology. 2012;22(1):53–66.

22. Tax CM, Jeurissen B, Vos SB, Viergever MA, Leemans A. Recursive calibration of the fiber response function for spherical deconvolution of diffusion MRI data. NeuroImage. 2014;86:67–80.

23. Tournier JD, Calamante F, Connelly A. Determination of the appropriate b value and number of gradient directions for high-angular-resolution diffusion-weighted imaging. NMR in biomedicine. 2013;26(12):1775–1786.

24. Mito R, Raffelt D, Dhollander T, et al. Fibre-specific white matter reductions in Alzheimer’s disease and mild cognitive impairment. Brain. 2018.

25. Jeurissen B, Tournier JD, Dhollander T, Connelly A, Sijbers J. Multi-tissue constrained spherical deconvolution for improved analysis of multi-shell diffusion MRI data. NeuroImage. 2014;103:411–426.

26. Schilling KG, Janve V, Gao Y, Stepniewska I, Landman BA, Anderson AW. Histological validation of diffusion MRI fiber orientation distributions and dispersion. NeuroImage. 2018;165:200–221.

27. Schilling K, Janve V, Gao Y, Stepniewska I, Landman BA, Anderson AW. Comparison of 3D orientation distribution functions measured with confocal microscopy and diffusion MRI. NeuroImage. 2016;129:185–197.

28. D’Arceuil HE, Westmoreland S, de Crespigny AJ. An approach to high resolution diffusion tensor imaging in fixed primate brain. NeuroImage. 2007;35(2):553–565.

29. Dyrby TB, Baare WF, Alexander DC, Jelsing J, Garde E, Sogaard LV. An ex vivo imaging pipeline for producing high-quality and high-resolution diffusion-weighted imaging datasets. Human brain mapping. 2011;32(4):544–563.

30. Schilling K, Gao Y, Stepniewska I, Choe AS, Landman BA, Anderson AW. Reproducibility and variation of diffusion measures in the squirrel monkey brain, in vivo and ex vivo. Magnetic resonance imaging. 2017;35:29–38.

31. Caruyer E, Lenglet C, Sapiro G, Deriche R. Design of multishell sampling schemes with uniform coverage in diffusion MRI. Magnetic resonance in medicine : official journal of the Society of Magnetic Resonance in Medicine / Society of Magnetic Resonance in Medicine. 2013;69(6):1534–1540.

32. Bigun J, Granlund GH. Optimal Orientation Detection of Linear Symmetry. Paper presented at: Proceedings of the IEEE First International Conference on Computer Vision; June, 1987.

33. Choe AS, Gao Y, Li X, Compton KB, Stepniewska I, Anderson AW. Accuracy of image registration between MRI and light microscopy in the ex vivo brain. Magnetic resonance imaging. 2011;29(5):683–692.

34. Rohde GK, Aldroubi A, Dawant BM. The adaptive bases algorithm for intensity-based nonrigid image registration. IEEE transactions on medical imaging. 2003;22(11):1470–1479.

35. Alexander DC, Pierpaoli C, Basser PJ, Gee JC. Spatial transformations of diffusion tensor magnetic resonance images. Medical Imaging, IEEE Transactions on. 2001;20(11):1131–1139.

36. Schilling KG, Gao Y, Stepniewska I, et al. The VALiDATe29 MRI Based Multi-Channel Atlas of the Squirrel Monkey Brain. Neuroinformatics. 2017.

37. Gao Y, Choe AS, Stepniewska I, Li X, Avison MJ, Anderson AW. Validation of DTI tractography-based measures of primary motor area connectivity in the squirrel monkey brain. PloS one. 2013;8(10):e75065.

38. Thomas C, Ye FQ, Irfanoglu MO, et al. Anatomical accuracy of brain connections derived from diffusion MRI tractography is inherently limited. Proceedings of the National Academy of Sciences of the United States of America. 2014;111(46):16574–16579.

39. Knosche TR, Anwander A, Liptrot M, Dyrby TB. Validation of tractography: Comparison with manganese tracing. Human brain mapping. 2015;36(10):4116–4134.

40. Dyrby TB, Sogaard LV, Parker GJ, et al. Validation of in vitro probabilistic tractography. NeuroImage. 2007;37(4):1267–1277.

41. Ferizi U, Schneider T, Tariq M, Wheeler-Kingshott CM, Zhang H, Alexander D. The Importance of Being Dispersed: A Ranking of Diffusion MRI Models for Fibre Dispersion Using In Vivo Human Brain Data. In: Mori K, Sakuma I, Sato Y, Barillot C, Navab N, eds. Medical Image Computing and Computer-Assisted Intervention – MICCAI 2013. Vol 8149: Springer Berlin Heidelberg; 2013:74–81.

42. Panagiotaki E, Schneider T, Siow B, Hall MG, Lythgoe MF, Alexander DC. Compartment models of the diffusion MR signal in brain white matter: a taxonomy and comparison. NeuroImage. 2012;59(3):2241–2254.

43. Schilling KG, Gao Y, Stepniewska I, Janve V, Landman BA, Anderson AW. Anatomical accuracy of standard-practice tractography algorithms in the motor system - A histological validation in the squirrel monkey brain. Magnetic resonance imaging. 2019;55:7–25.

44. Maier-Hein KH, Neher PF, Houde JC, et al. The challenge of mapping the human connectome based on diffusion tractography. Nat Commun. 2017;8(1):1349.

45. Stepniewska I, Preuss TM, Kaas JH. Thalamic connections of the primary motor cortex (M1) of owl monkeys. J Comp Neurol. 1994;349(4):558–582.

46. Stepniewska I, Preuss TM, Kaas JH. Architectonics, somatotopic organization, and ipsilateral cortical connections of the primary motor area (M1) of owl monkeys. J Comp Neurol. 1993;330(2):238–271.

